# Bedshift: perturbation of genomic interval sets

**DOI:** 10.1101/2020.11.11.378554

**Authors:** Aaron Gu, Hyun Jae Cho, Nathan C. Sheffield

## Abstract

Functional genomics experiments, like ChIP-Seq or ATAC-Seq, produce results that are summarized as a region set. Many tools have been developed to analyze region sets, including computing similarity metrics to compare them. However, there is no way to objectively evaluate the effectiveness of region set similarity metrics. In this paper we present *Bedshift*, a command-line tool and Python API to generate new BED files by making random perturbations to an original BED file. Perturbed files have known similarity to the original file and are therefore useful to benchmark similarity metrics. To demonstrate, we used Bedshift to create an evaluation dataset of hundreds of perturbed files generated by shifting, adding, and dropping regions from a reference BED file. Then, we compared four similarity metrics: Jaccard score, coverage score, Euclidean distance, and cosine similarity. Our results highlight differences in behavior among these metrics, such as that Jaccard score is most sensitive to added or dropped regions, while coverage score is most sensitive to shifted regions. Together, we show that Bedshift is a useful tool for creating randomized region sets for a variety of uses.

**Availability:** BSD2-licensed source code and documentation can be found at https://bedshift.databio.org.

## Introduction

In the past few years, projects such as ENCODE (Encyclopedia of DNA Elements) and IHEC (International Human Epigenome Consortium) have established large catalogs of genomic features, including regulatory regions, transcription factor binding sites, and SNPs (Dozmorov, 2017). These data can be summarized into region sets, often stored in BED file format containing genomic regions represented by a chromosome number, start position, stop position, and optional metadata. Increasingly, computational tools are being developed to produce and consume BED files (Zhou *et al*., 2020). Early studies used interval analysis to study regulatory elements for biological conclusions (Zhang *et al*., 2007; Wederell *et al*., 2008; Chen *et al*., 2008; Johnson *et al*., 2007; Fu *et al*., 2008; Cuddapah *et al*., 2009), and region sets are of particular interest in epigenomics, where hundreds of thousands of cell-type specific elements have been shown to play an important part in gene regulation (Song *et al*., 2011; Sheffield and Furey, 2012; Thurman *et al*., 2012).

Among the many applications of region sets, interest has grown in methods to compare region sets with one another (Kanduri *et al*., 2018). New genomic regions produced from experiments can be associated with established genomic regions using co-occurrence, under the assumption that region sets with many overlapping regions may reflect biological relationships (Dozmorov, 2017). There are many methods to evaluate the similarity of two region sets, which have been under development for more than a decade (Fu and Adryan, 2009; Zhang *et al*., 2007; Chen *et al*., 2008; Huen and Russell, 2010; Carstensen *et al*., 2010; Chikina and Troyanskaya, 2012; Heger *et al*., 2013; Khushi *et al*., 2014; Sarmashghi and Bafna, 2019; Ferré *et al*., 2019; Feng *et al*., 2020). One general tool that provides the user with multiple results is the GSuite Hyperbrowser, which includes the most similar region sets, unique region sets, and how the co-occurrence counts change along the genome (Simovski *et al*., 2018). Some tools use a statistical test to measure the significance of the co-occurrence.

For example, GenomeRunner (Dozmorov *et al*., 2016), LOLA (Sheffield and Bock, 2016; Nagraj *et al*., 2018), GIGGLE (Layer *et al*., 2018), and IGD (Feng and Sheffield, 2020) take BED files as input and compute region overlap counts, followed by a Fisher’s exact test to produce a similarity score or ranking of most similar files. Other tools, such as regioneR and ChIP-Seeker, use permutation or sampling of regions or random background region set to calculate the probability of observing more extreme overlap between it and the provided data (Yu *et al*., 2015; Gel *et al*., 2015). Another approach is to examine distribution along the genome such as the approach taken by GenometriCorr (Favorov *et al*., 2012). There is therefore a wide variety of methods and tools to assess relationships among region sets (Kanduri *et al*., 2018; Simovski *et al*., 2018). These tools are similar in their attempt to compare region set data, but have subtle differences in the goal, data used, and approach of comparison.

Here, we provide a conceptual framework upon which similarities among region sets may be evaluated. We do this by simulating perturbations of region sets, allowing us to construct ground truth results between two region sets. We introduce Bedshift, a command line interface and Python package that provides users the ability to create new BED files based on random modifications to an original BED file. A user can specify what percentage of regions they want to shift, drop, add, cut, and merge. Users may also specify for each perturbation subsets of regions to perturb using separate selector region sets. The most similar existing tool to Bedshift is a function in the BEDTools suite called `shuffleBed` (Quinlan, 2014). This function randomly permutes the regions inside a BED file, moving them to different locations in the genome while preserving their length, which is useful for generating background or randomized region sets. Bedshift provides control over the type, magnitude, and combinations of perturbation, and makes it easy to produce many replicates, making it suitable for more complex perturbations and to test how similarity metrics behave with different types and levels of perturbation.

Bedshift produces reference files that are useful for many downstream tasks, including as controls for experimental region set data, as a randomized background of region data, as test data for a new tool, or to test similarity metrics. In this paper, to demonstrate one use case, we applied Bedshift to evaluate region set similarity metrics. We created an evaluation set of thousands of files with controlled levels of divergence to an original file, and then compared different similarity metrics to see how scores vary as the type and level of perturbation changes. This study reveals that similarity metrics vary in sensitivity to different types of perturbation, and that for universe-based metrics, the choice of universe is a critical experimental decision.

## Results

### Overview of Bedshift

Bedshift is available as a command line interface as well as a Python package. Documentation with guides on common use cases is available at bedshift.databio.org. Bedshift perturbs regions in a region set with 5 possible operations: shift, add, cut, merge, and drop. The operations can be specified one at a time or all in one command, in which case Bedshift will run them in the order of shift, add, cut, merge, and then drop. The number of regions perturbed is set as a proportion of the total number of regions in the region set. For example, if given a BED file with 1000 regions, the operation `bedshift -b example.bed -a 0.2 -s 0.4` would first shift 400 of the regions (40%), then add 200 new regions (20%).

The shift operation will shift the start and end position of a region by a random value based on a normal distribution specified by the user using the `--shiftmean` and `--shiftstdev` options. In conrast to BEDTools, Bedshift does not shift regions to a completely new location on the genome, but upstream or downstream by a small, random number, placing them near their original location. The add operation will create randomly generated regions on any chromosome with a length based on a normal distribution specified by the user using the `--addmean` and `--addstdev` options. The drop operation will randomly delete regions from the region set. The cut operation will split a region into two new regions, with the split position in the region being randomly determined. Finally, the merge operation will merge two adjacent regions, potentially creating very large regions.

In addition to these five operations, we have added numerous features to give the user more configurability. The `--addfile`, `--dropfile`, and `--shiftfile` options allow users to input a file from which regions are selected to be added, dropped, or shifted. This feature makes Bedshift able to configure perturbation type and level to specific region types, such as introns, exons, promoters, or enhancers. To facilitate dataset generation, the `--repeat` option makes it easy to create many replicates of the perturbation with a single command.

Users may specify perturbations on the command line, from within Python, or using a YAML configuration file with the `--yaml-config` option. This yaml configuration file allows users to specify the order or perturbations and construct arbitrary complex combinations, which also make it possible to construct highly realistic biological scenarios, such as dropping only a subset of promoter elements or adding from a prespecified list of enhancer elements. In the documentation, we provide scripts that show how bedshift can be used to create thousands of perturbed files for dozens of different parameter sets easily with a few commands on the command line.

### Simulation study and evaluation approach

To test Bedshift and demonstrate how it can be used to evaluate similarity measures of region sets, we selected one input file from ENCODE (ENCFF549PGC)(Moore *et al*., 2020) and created an evaluation set of perturbed BED files for *shift, drop*, and *add*, with 3 levels and 10 replicates for each perturbation. Our parameter values included a low, medium, and high degree for each perturbation type, and visual inspection of region sets allowed us to tune the parameters to a biologically relevant range (Figure 1B; Figure S1). We expected that similarity scores would reflect this known degree of perturbation. We used the `--addfile` feature of Bedshift to add regions from a “universe” of possible regions to include, instead of completely random regions. Our primary universe is unified set of regulatory elements from the SCREEN database of the ENCODE project (See Methods) (Moore *et al*., 2020).

**Figure 1:**
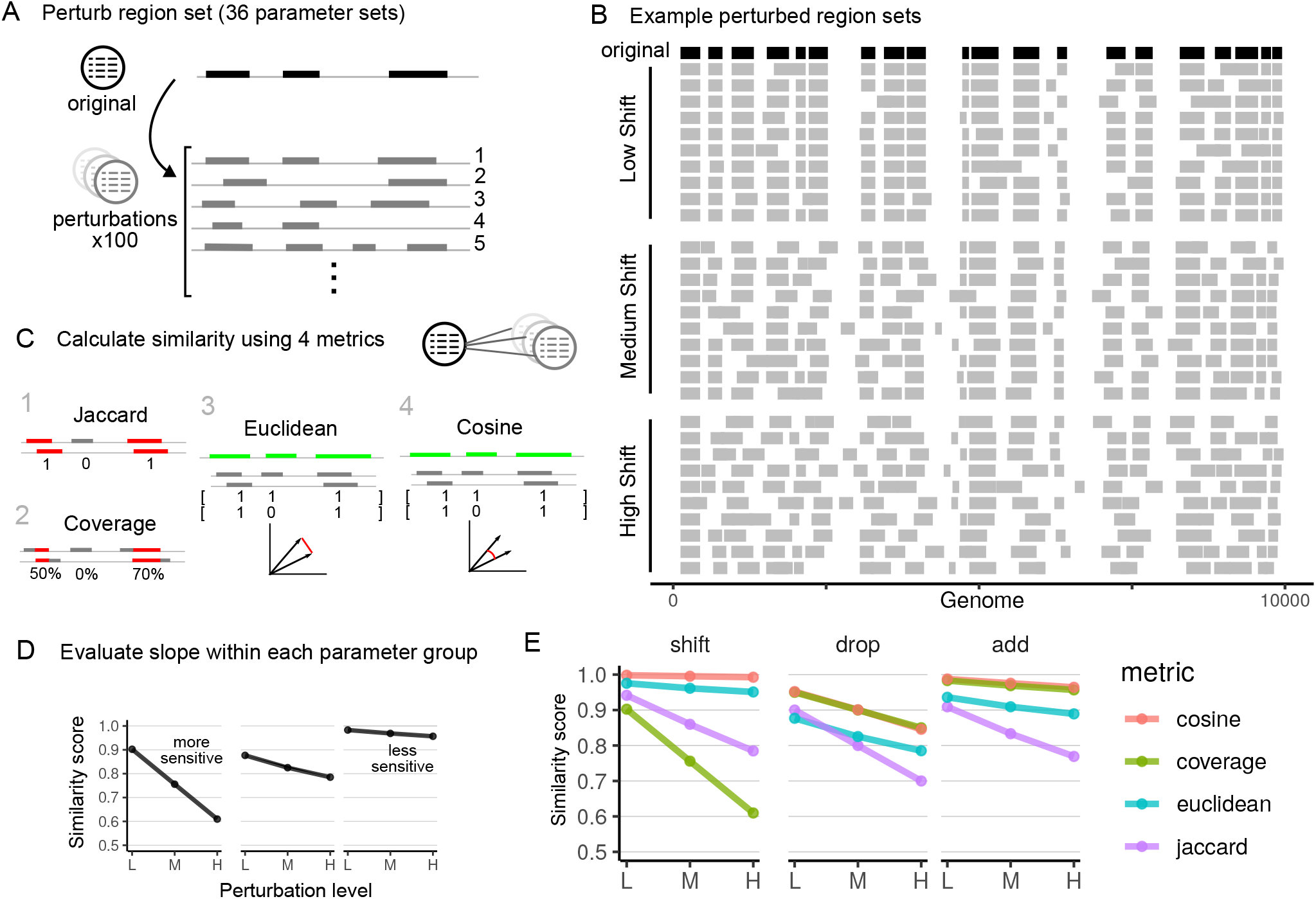
Overview of simulation study and comparison of four similarity metrics. A) One BED file was used to create 3600 perturbed files, 100 repetitions for each of 36 different combinations of add, drop, and shift perturbations. B) Examples of a demo region set that has been shifted to different degrees. C) We calculated four similarity metrics between the original file and the perturbed files. The greater the similarity score decrease, the more sensitive the metric is. **TODO** change slope in figure D) Within each parameter group, as the perturbation increases, the similarity score decreases. E) Results for shift, drop, and add-only perturbations. L-Low; M-Moderate; H-High perturbation.

To extend this basic experiment, we did three additional, extended experiments: First, to test how the metrics behave in combinations of perturbations, we expanded the experiment to include pairwise combinations of each level and perturbation, resulting in 360 BED files made from 36 different Bedshift parameter sets (Supplemental Materials Table S1), such as adding and shifting, or dropping and shifting, repeating each combination 10 times (Figure 1A). Second, we repeated this combinatorial study on 3 separate input files to test how the input file affects the metrics. Finally, we repeated this study using the original file, but with 3 additional universes, to test how changing the universe affects the metrics. Additional universes are specific subsets of the primary universe from SCREEN: CTCF sites, promoter-like sequences (PLS), or DNase-H3K4me3 sites (See Methods).

After simulating perturbed region sets to known parameters, we sought to evaluate the performance of different measures of region set similarity. For each pairwise comparison of original to perturbed BED file, we computed four similarity metrics: Jaccard score, coverage score, Euclidean distance, and cosine similarity (Figure 1B). The Jaccard score and coverage metrics were chosen based on their common usage in other similarity scoring methods (Kanduri *et al*., 2018). The Euclidean distance and cosine similarity metrics are a simple vector-based approach computed on binary vectors with each element reflecting presence or absence of a region in the universe. We focused in this study on metrics useful for measuring the level of difference between two very similar region sets, as opposed to other common tools (such as the Fisher’s Exact Test) which can be used to test the hypothesis that two sets are independent. To evaluate the metrics, we compared each measure for its ability to reflect Bedshift perturbations (Figure 1D).

### Experiment 1: Evaluating individual perturbations

#### Shift

As the percentage of *shift* was increased from 20% to 50% to 80%, the similarity score decreased for all four scoring methods.

The score with the greatest amount of decrease was the coverage score, which can be explained by how the coverage score measures the percentage of individual regions that overlap with other regions, and is therefore affected by any shift. In contrast, the other measures are based only on overlap counts, which will only change if the shift is substantial enough to eliminate overlap.

#### Drop

When measuring the drop perturbation, which increased from 10% to 20% to 30%, the Jaccard score had the greatest similarity score decrease (Figure 1D). In fact, the way in which the Jaccard score is calculated makes it so that it measures the exact percentage of regions dropped. Shown in the graph, the Jaccard score decreases perfectly from 90% to 80% to 70% as drop increases from 10% to 20% to 30%. The other three metrics displayed levels of decrease which were smaller than the Jaccard score decrease. This indicates that the simple Jaccard overlap counting method was the most sensitive to dropped regions.

#### Add

For the *add* perturbation, the Jaccard score again had the lowest similarity scores and steepest decline as the *add* percentage increased from 10% to 20% to 30% (Figure 1D). However, unlike in the *drop* perturbation, the Jaccard score does not perfectly measure the percentage of added regions. This is due to added regions having the probability of overlapping existing regions, which would thus not be recognized as a newly introduced region. Interestingly, not only was the Jaccard score less sensitive, but we also found that the other three metrics were less sensitive to adding regions than dropping regions. We expected sensitivity to adding to resemble sensitivity to dropping, because the perturbations are complementary. We explore this further in the next section.

#### Sensitivity of dropping vs adding

We were initially surprised that all metrics were more sensitive to dropping than adding. After employing a hypothetical overlap counting example, we can see why that happens for the Jaccard score: Given a set of 4 regions, if a non-overlapping region is added, the score would decrease by 25% to 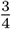. On the other hand, if a region is dropped, then the score decreases by 33% to 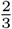. This provides a theoretical explanation of the observation that the Jaccard score is more sensitive to drop than add. Remarkably, our perturbation results were able to capture this nuance. Similarly, the coverage score is less sensitive to add than drop, which can be explained similarly: adding a region would decrease the score by 12.5% to 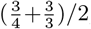, and dropping a region would decrease the score by 16.7% to 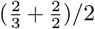. Thus, dropping regions still decreases the coverage score more than adding.

### Experiment 2: Evaluating combinatorial perturbations

To extend our results, we next examined the sensitivity when combinations of perturbations were used. We used Bedshift to create perturbed files with each pairwise combination of parameters, at each level (Table S1). This resulted in 36 parameter sets: 9 represent the 3 individual perturbations at 3 levels each discussed previously, and then 27 sets represent each pairwise combination of 3 perturbations at 3 levels. We grouped these results into 9 scenarios, 3 of which represent individual perturbations and 6 of which represent each pairwise comparison of perturbation (Figure 2A). For example, in *Scenario 2*, we plotted the decrease of the similarity score as the *add* perturbation is increased, with the *drop* perturbation held constant. For each pairwise scenario, we used 3 different levels (high, moderate, and low) of one type of perturbation in combination with another perturbation held constant, resulting in 3 plots per pairwise scenario. To summarize these results, we computed the decrease across three levels of perturbation increase (Figure 2B) in each line plot. As a further summary, we created a heat map of each scores’ sensitivity to the three perturbations we tested (Figure 2C).

**Figure 2:**
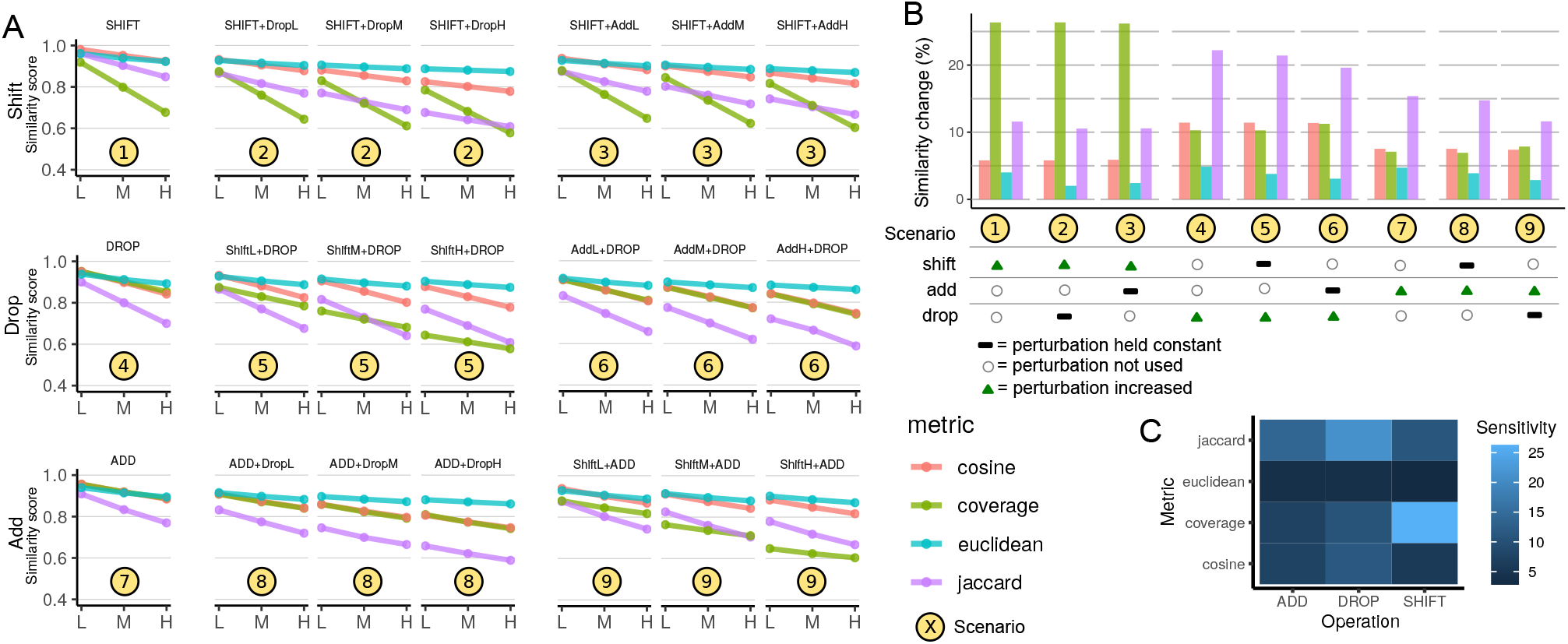
Bedshift and similarity score results. A) Similarity scores for shift, drop, and add perturbations, along with their pairwise combinations. B) Summary of similarity score change for different perturbation levels measured. For a given scenario, each perturbation is classified as either “increasing”, “not used”, or “held constant”. To condense the information from panel A, the amount of decrease shown for each Scenario is the average across all three levels in the corresponding Scenario in panel A. C) Summary of the sensitivity of each metric to the three different perturbations.

### Multiple perturbations decrease similarity score and preserve sensitivity

OUr results show that the similarity scores for combinatorial perturbations are lower than each perturbation individually, showing that the similarity scores accurately recognize that more perturbation, even of different types, leads to lower similarity (Figure 2A). Furthermore, the overall sensitivity trends remain intact (Figure 2B). For instance, the coverage score is clearly still most sensitive to *shift*, even in the presence of *add* and *drop*. Similarly, the Jaccard score remains most sensitive to *drop*, and, to a lesser degree, to *add* in the combinatorial analysis. Furthermore, all metrics remain more sensitive to *drop* than to *add* (Figure 2B). This indicates that these metrics are robust and detect changes that are compounded on each other.

### Euclidean distance is the least sensitive overall

This analysis also shows that across all individual and combinatorial perturbations, our Euclidean distance metric is the least sensitive to changes (Figure 2C). This result indicates that these metrics could be useful for different purposes, with Euclidean distance as implemented here more likely to be useful for more distant relationships among region sets. Interestingly, the cosine similarity and coverage scores behave almost identically to the *add, drop*, and *add* + *drop* scenarios, but differ dramatically when *shift* is included, due to the increased sensitivity of coverage to *shift*.

### Experiment 3: Testing across input files

To ensure that the results are not specific to input file, we ran the experiment again on two additional files. The original file contained CTCF data with an average region length of 301 bp, and the other two files contain DNA methylation data and DNAse-seq data, with average region length of 1161 bp and 150 bp, respectively. These therefore reflect a variety of data types and region sizes. The three files experiment shows that, in general, the original analysis results hold across all three input files: coverage is most sensitive to *shift*; jaccard is most sensitive to *add* and *drop*;

Euclidean distance is least sensitive overall; *etc*. (Figure S2). However, this analysis also reveals an interesting observation that the metrics do behave differently for the different files. Most pronounced, we observe that different region length in the two new files caused results to vary in all of the shift perturbation combinations, especially for the coverage score (Figure S2, Scenario 1). The file with the highest similarity score decrease was the file with the smallest average region length, while the file with larger regions had a less pronounced sensitivity to *shift*. This reflects the constant shift distance across the files, so the file with the smallest region length would be the most likely to have regions shifted further from their original locations. In conclusion, specific similarity results are clearly affected by input file, but general sensitivity trends among metrics hold across input files.

### Experiment 4: Testing across universes

In addition to the multiple file analysis, we also wanted to see if the universe choice would impact the results. We chose to use three subsets of the SCREEN universe: CTCF sites only, promoter-like sequences (PLS), or DNase-H3K4me3 sites, and re-ran the analysis using the original file, but switching the universe (Figure S3). We observed that the coverage and Jaccard scores were invariant across the three universes under the same operations, whereas Euclidean distance and cosine similarity varied significantly. This is expected, as the coverage and the Jaccard scores are not vector-based similarity measures, while Euclidean distance and cosine similarity are, and therefore depend on the chosen universe. If the universe does not encapsulate the regions in the file we are perturbing, then the Euclidean distance and cosine similarity score will be less accurate. In addition, we noticed that as the size of the universes decreased, Euclidean distance and cosine similarity became more sensitive. This can be explained by the proportion of vector components that change. When the dimension of a vector is smaller, one component of the vector changing due to an add, drop, or shift will cause the vector to change proportionally more.

In Scenario 6, we observe an interesting anomaly, that for the Euclidean distance, the slope turns positive, indicating that when adding regions is held constant, dropping more regions actually *increases* similarity between the files. This counterintuitive result can be explained by the possibility of dropping regions that were just added. In most cases, this is not a problem because the probability of dropping regions that were just added is low; however, in very specific situations, such as this particular scenario in our study, this effect can occur. The effect would be most pronounced when universes are very different from the query file, because the *add* operation is adding regions unlikely to be in the original file, making it more likely that drop will drop something that’s different from the original file, increasing similarity. Also, it makes sense that this is more pronounced when *add* is higher, because it increases probability of dropping a region that is different. Further, we can explain that it only occurs for Euclidean distance, not for cosine distance, because for widely divergent universes, when adding lots of new unrelated dimensions, cosine distance is unaffected, as the angle between two vectors is only calculated on the basis of dimensions present; in contrast, Euclidean distance explodes. This result emphasizes the importance of choosing an appropriate, fitting universe for vector-based approaches; if the universe is not a good reflection of the underlying data, then distance metrics may potentially behave counterintuitively, particularly for the Euclidean distance

### Future Development

We have shown a use case for Bedshift in investigating how region set similarity metrics are affected by different types of differences. There are several ways Bedshift could be extended to address additional questions. First, Bedshift does not consider strandedness in perturbations, but adding strand-aware perturbations would allow testing how strand-aware similarity metrics behave. There could also be a new perturbation that flips the strand. Another extension is a perturbation similar to BedTools shuffle that moves regions to completely new locations in the genome, but does not change their length. Finally, we are working on ways to increase efficiency of the shift operation, which currently is the slowest operation because it iterates over and edits each region.

## Conclusion

In this paper we present Bedshift, a new tool to help researchers evaluate the effectiveness of region set similarity metrics. Similarity scoring metrics and tools are becoming increasingly common, and it is important to know how each tool performs on different datasets. Bedshift is a way to generate new BED files with perturbations such as shifted regions, added regions, dropped regions, and more.

Our results provide an initial analysis to compare different similarity scoring metrics. In our study design, we considered scores that measure differences among similar region sets, as opposed to scores that test a hypothesis that region sets are independent. We uncovered interesting information about the relative performance of these metrics. The key conclusion is that similarity scores have unique sensitivities to types of perturbation. One key caution is that a metric that is more sensitive will more quickly reach a saturation point; at this stage, the metric becomes unreliable. In our analysis, we did not identify a global “best” metric, but each metric is likely to be more appropriate depending on use case. For example, Euclidean distance and cosine similarity were generally the least sensitive overall and would therefore be more useful for measuring similarities between distant region sets. Overall, the Jaccard score seemed to be the most sensitive, and may therefore be most useful for highly similar region sets. It also showed consistency in the slope decrease across multiple levels of perturbation. The coverage score would be most appropriate for detecting slight shifts. Discovering the performance of these different similarity metrics showcases a powerful use for Bedshift, as it has allowed us to discover pros and cons of different similarity metrics.

Our analysis also leads to several directions for future work. First, it is possible that the universe-based measures would behave differently depending on the universe used to construct the vectors. More work needs to be done to explore optimal ways for constructing universes, which would benefit vector-based similarity metrics. Second, it will be interesting to explore the behavior of new similarity metrics in comparison to current similarity metrics. Finally, similarity scoring methods could be combined for increased confidence in results. Bedshift will be a helpful tool going forward as we develop and evaluate new ways to measure similarity of region sets.

## Availability of Data and Materials

For this paper, we used bedshift version 1.1.0. Bedshift is licensed under BSD-2, and can be downloaded from GitHub: https://github.com/databio/bedshift/ or from the Python Package Index (https://pypi.org/project/bedshift/). Complete documentation can be found at bedshift.databio.org, including repositories with code to reproduce the all experiments in this paper. All datasets used are publicly available with download instructions included in the documentation.

## Acknowledgments

We would like to thank Erfaneh Gharavi and Guangtao Zheng for helpful comments.

## Methods

### Data set

The 3 query files used as the original file, which is then perturbed with Bedshift are all from the ENCODE consortium: 1) CTCF TF ChIP-seq on human HCT116 (primary file, ENCFF549PGC); 2) K4me3 Histone ChIP-seq on human GM12864 (ENCFF749NUK); and 3) DNase-seq on human stromal cell of bone marrow (ENCFF409URA).

The 4 universe region sets used in this analysis are from the SCREEN ENCODE database: 1) GRCh38-ccREs (Primary universe); 2) DNase-H3K4me3; 3) CTCF-only; and 4) PLS.

### Bedshift operations

The order of operations is shift, add, cut, merge, and drop. From the command line interface, a call to `bedshift` will use the `all_perturbations` function to run up to all 5 perturbations, then output the file either in a user specified location using the `--outputfile` option, or in the same directory with the original filename prepended with `bedshifted_`. From the Python API, the user has more flexibility to call the perturbations individually, or use the same `all_perturbations` function. Bedshift stores the state of the BED file in a Pandas dataframe, and each perturbation operates on the result of the previous one, which is why the order is important. When using the Python API, the state of the BED file can be reset to its original state using the `reset_bed` () method.

### Shift

Using the rate parameter as a proportion of the total number of regions in the BED file, a subset of regions to shift is chosen from the BED file. The start and end position of each of these regions is adjusted by the same distance. The distance is chosen from a normal distribution (`--shiftmean, --shiftstdev`), which defaults to N(0, 150). In order to use shift, a .fasta file containing chromosome lengths must be provided with the `--chromosome-lengths` option, to ensure that regions are not shifted off the ends of chromosomes.

### Shift From File

Shift from file uses a file specified through `--shiftfile` to determine which regions to shift. If `pyranges` tool is available on the user’s machine, then `--shiftfile` will consider only intersecting regions as candidates to be shifted. Otherwise, only regions that are exact matches will be candidates for shifting.

### Add

The number of regions to add is determined from the rate parameter as a proportion of the total number of regions in the BED file. For each added region, first a chromosome is chosen with proportional odds to the length of each chromosome; then a start position is chosen anywhere along the chromosome; then a length is computed based on a normal distribution defaulting to N(320, 20) (`--addmean, --addstdev`) and added to the start position to arrive at the end position. In order to use add, a .fasta file containing chromosome lengths must be provided with the `--chromosome-lengths` option.

### Add From File

Instead of adding randomly generated regions, the user can specify a file to the `--addfile` option which contains candidate regions to add. From these regions, a number of them is chosen based on the rate parameter and added to the file.

### Add in Valid Regions

Another way to add regions is to specify a file with valid regions where random new regions can be added. The option is called `--valid-regions`. The user would use this option instead of `--addfile` if they wanted to add more random regions instead of specific regions from a file, but wanted to restrict these to certain valid areas of the genome. The valid regions file could contain very large areas of the genome such as introns or promoters. Random region generation works the same as the basic add operation, but restricts the chromosomes and regions to the ones specified in the `--valid-regions` file.

### Cut

Using the rate parameter as a proportion of the total number of regions, the regions to cut are determined. For each of these regions, the cut is made at the midpoint of the start and end position, creating two new regions. The original region is dropped from the BED file.

### Merge

Using the rate parameter as a proportion of the total number of regions, the regions to merge are determined. For each of these regions, they are merged with the next subsequent region in the BED file if they are both on the same chromosome by taking the start position of the first region and the end position of the second region. We recognize that this method has the potential to create very large regions.

### Drop

Using the rate parameter as a proportion of the total number of regions, the regions to drop are determined. They are simply removed from the BED file.

### Drop From File

Similar to add from file and shift from file, drop from file uses a file specified through `--dropfile` to determine which regions to drop. If `pyranges` tool is available on the user’s machine, then `--dropfile` will consider only intersecting regions as candidates to be dropped. If those tools are not available, then only regions that are exact matches will be candidates.

### Seed

Sometimes, users may wish their Bedshift perturbations to be identically reproducible. Assuming every other operation remains constant, setting the same integer-valued seed through `--seed` will allow Bedshift to produce identical perturbations.

### Bedshift file generation

We leverage the Looper tool (http://looper.databio.org) to create hundreds of perturbation replicates using the same 4 columns (sample_name, shift, add, and drop) as Supplemental Table 1. The normal distributions used in shift and add are the default parameters. Each sample was run by looper using a Bedshift command with the specified shift, add, and drop. Details and actual scripts used in the analysis can be found in the documentation for bedshift.

### Jaccard score

The first metric was the Jaccard score based on overlapping regions between two BED files, computed by the formula

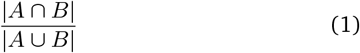

where *A* and *B* are the two BED files. Any region with at least 1 base pair overlap was counted in the total number of overlaps. Overlaps were computed using Augmented Interval List (AIList) (Feng *et al*., 2019).

### Coverage score

The BedTools coverage function was used, which takes in two BED files and uses the first one as the reference region set to determine coverage for each region in the second BED file (Quinlan, 2014). A normalization technique was applied to assign coverage scores to every region in both files. First, BedTools coverage (in the Python wrapper PyBedTools) was run with the perturbed file as the first argument and the original file as the second. No additional arguments were provided other than the two files. Then the files were passed as arguments to the coverage tool in the opposite order, in order to account for coverage in regions across both files. This produced a coverage score between 0 and 1 for each region in both the original and the perturbed file. To get the final similarity score, the mean was taken of coverage values for every region in both files:

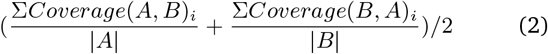

where *A* and *B* are the two BED files and *Coverage* is the BedTools coverage score.

### Euclidean distance

In vector-based similarity methods, a standard vocabulary was needed to represent each region as a position in the vector. To do this, a “vocabulary”, or a universe, was used. For our primary analyses, we used the general set of regulatory elements from the SCREEN database ((Moore *et al*., 2020)). We also tested other universes in the universe experiment. When casting new files into the universe to create a vector, if a region in the new file overlapped with a universe region, then that index in the vector was set to 1. Therefore, each BED file was represented by a vector of 0’s and 1’s. The Euclidean distance is defined as

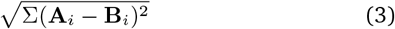

where **A** and **B** are the two vectors representing the BED files. A normalized Euclidean distance was calculated by dividing by the maximum distance of two vectors (a vector of all 0’s and a vector of all 1’s), which was 518.02. That value was subtracted from 1, because a smaller normalized distance indicates a higher similarity.

### Cosine similarity

The same vectorization technique and vectors used for the Euclidean distance metric were also used for the cosine similarity analysis. The cosine similarity is defined as

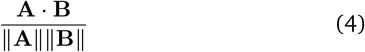

where **A** and **B** are the two vectors representing the BED files. In vector space, the closer two vectors are, the closer their cosine is to 0. Thus, the resulting cosine score was subtracted from 1 to get the final similarity score.

### Similarity score change

The change in similarity was measured by taking the difference between the highest and lowest score in the perturbation level (for example, the score difference between add 0.1 and add 0.3 using the Jaccard score was 0.15). For groups of 9 perturbation parameters, such as increasing shift from 0.1 to 0.3 while holding add constant at 0.1, 0.2, and 0.3, the three scores from the levels of add were averaged.

## Supplemental Materials

**Table S1:** Parameter combinations used in the analysis.

| parameter set | add | drop | shift |
| --- | --- | --- | --- |
| add1 | 0.1 | 0.0 | 0.0 |
| add2 | 0.2 | 0.0 | 0.0 |
| add3 | 0.3 | 0.0 | 0.0 |
| drop1 | 0.0 | 0.1 | 0.0 |
| drop2 | 0.0 | 0.2 | 0.0 |
| drop3 | 0.0 | 0.3 | 0.0 |
| shift1 | 0.0 | 0.0 | 0.2 |
| shift2 | 0.0 | 0.0 | 0.5 |
| shift3 | 0.0 | 0.0 | 0.8 |
| add_drop1 | 0.1 | 0.1 | 0.0 |
| add_drop2 | 0.1 | 0.2 | 0.0 |
| add_drop3 | 0.1 | 0.3 | 0.0 |
| add_drop4 | 0.2 | 0.1 | 0.0 |
| add_drop5 | 0.2 | 0.2 | 0.0 |
| add_drop6 | 0.2 | 0.3 | 0.0 |
| add_drop7 | 0.3 | 0.1 | 0.0 |
| add_drop8 | 0.3 | 0.2 | 0.0 |
| add_drop9 | 0.3 | 0.3 | 0.0 |
| shift_drop1 | 0.0 | 0.1 | 0.2 |
| shift_drop2 | 0.0 | 0.1 | 0.5 |
| shift_drop3 | 0.0 | 0.1 | 0.8 |
| shift_drop4 | 0.0 | 0.2 | 0.2 |
| shift_drop5 | 0.0 | 0.2 | 0.5 |
| shift_drop6 | 0.0 | 0.2 | 0.8 |
| shift_drop7 | 0.0 | 0.3 | 0.2 |
| shift_drop8 | 0.0 | 0.3 | 0.5 |
| shift_drop9 | 0.0 | 0.3 | 0.8 |
| add_shift1 | 0.1 | 0.0 | 0.2 |
| add_shift2 | 0.1 | 0.0 | 0.5 |
| add_shift3 | 0.1 | 0.0 | 0.8 |
| add_shift4 | 0.2 | 0.0 | 0.2 |
| add_shift5 | 0.2 | 0.0 | 0.5 |
| add_shift6 | 0.2 | 0.0 | 0.8 |
| add_shift7 | 0.3 | 0.0 | 0.2 |
| add_shift8 | 0.3 | 0.0 | 0.5 |
| add_shift9 | 0.3 | 0.0 | 0.8 |

**Figure S1:**
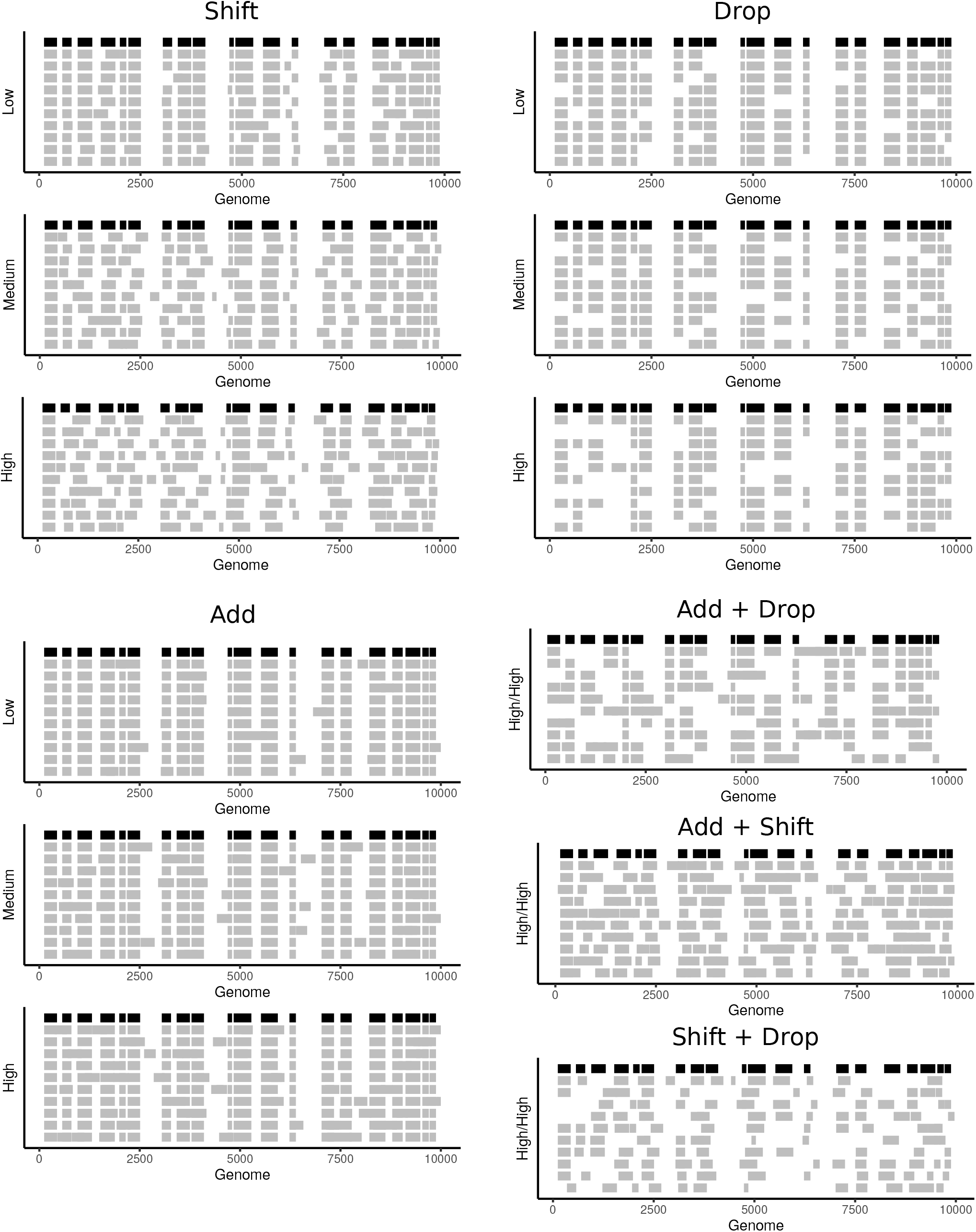
Demo visualization of perturbations. Original file is shown in black, randomizations in gray. The vertical axis label sdepict perturbation degree, corresponding to parameter values in the parameter table. Shift plots are reproduced here from figure 1 for comparison. For combinatorial perturbations, only the maximum perturbation is shown.

**Figure S2:**
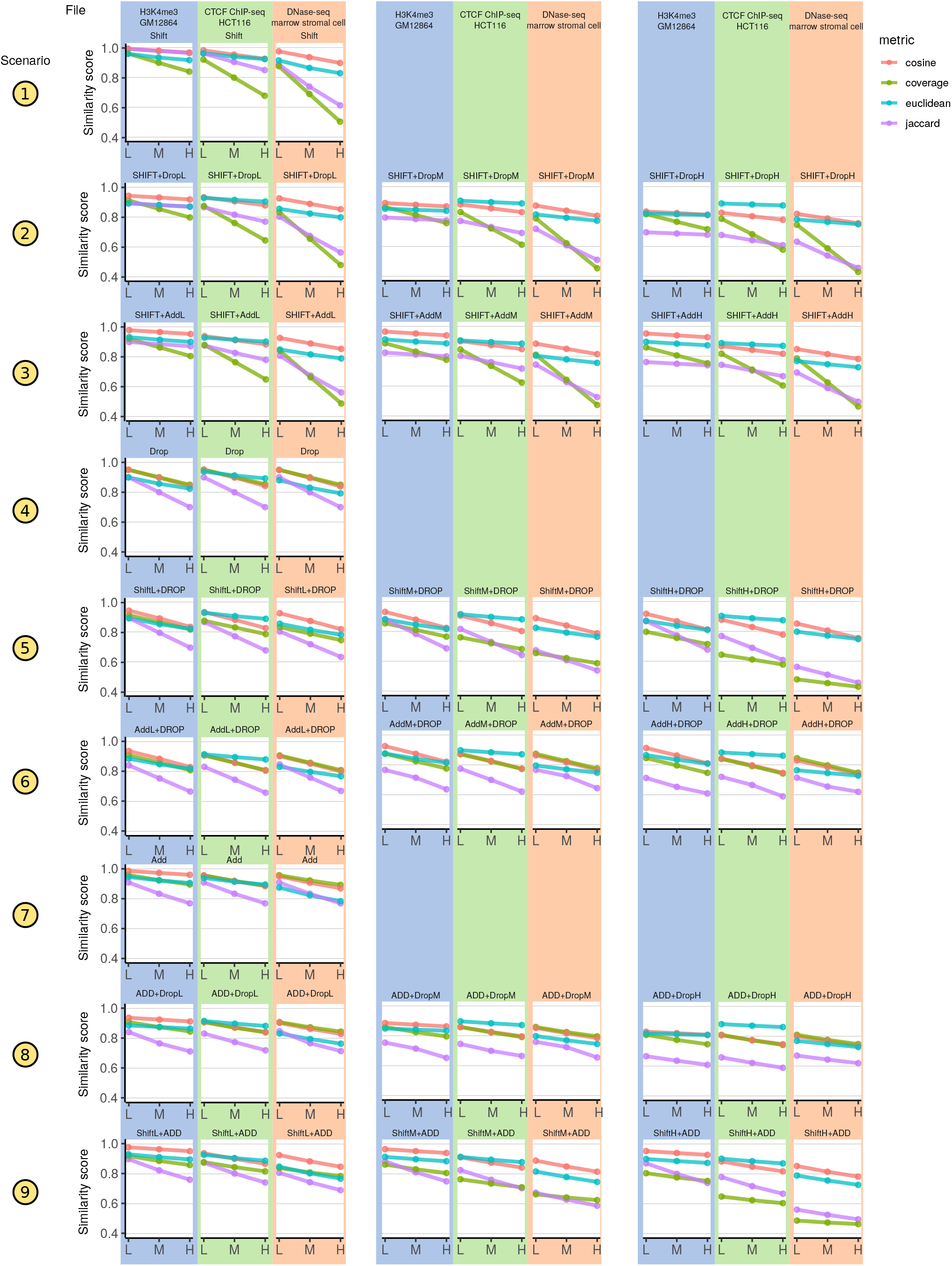
Detailed results showing how metrics vary by query file. Universe is held constant.

**Figure S3:**
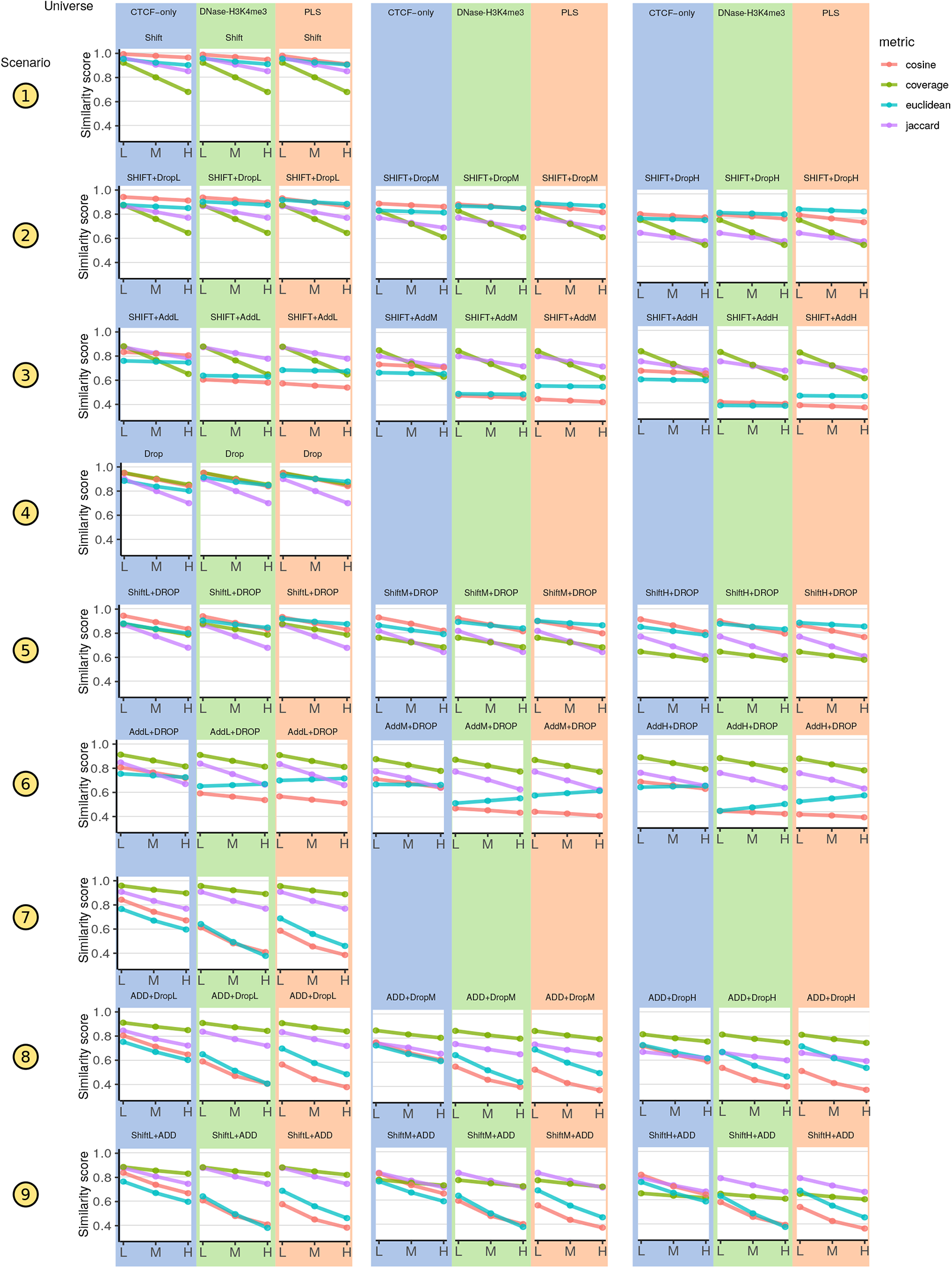
Detailed results showing how metrics vary by universe. Query file is held constant, and is the same as in the original study.

## References

Carstensen, L. et al. (2010) Multivariate hawkes process models of the occurrence of regulatory elements. BMC Bioinformatics, 11, 456.

Chen, X. et al. (2008) Integration of external signaling pathways with the core transcriptional network in embryonic stem cells. Cell, 133, 1106–1117.

Chikina, M.D. and Troyanskaya, O.G. (2012) An effective statistical evaluation of ChIPseq dataset similarity. Bioinformatics, 28, 607–613.

Cuddapah, S. et al. (2009) Global analysis of the insulator binding protein CTCF in chromatin barrier regions reveals demarcation of active and repressive domains. Genome Res., 19, 24–32.

Dozmorov, M.G. (2017) Epigenomic annotation-based interpretation of genomic data: From enrichment analysis to machine learning. Bioinformatics, 33, 3323–3330.

Dozmorov, M.G. et al. (2016) GenomeRunner web server: Regulatory similarity and differences define the functional impact of SNP sets. Bioinformatics, 32, 2256–2263.

Favorov, A. et al. (2012) Exploring massive, genome scale datasets with the GenometriCorr package. PLoS Computational Biology, 8, e1002529.

Feng, J. et al. (2019) Augmented interval list: A novel data structure for efficient genomic interval search. Bioinformatics.

Feng, J. and Sheffield, N.C. (2020) IGD: High-performance search for large-scale genomic interval datasets. Bioinformatics.

Feng, S.C. et al. (2020) Seqpare: A self-consistent metric of similarity between genomic interval sets. F1000Research, 9, 581.

Ferré, Q. et al. (2019) OLOGRAM: Determining significance of total overlap length between genomic regions sets. Bioinformatics.

Fu, A.Q. and Adryan, B. (2009) Scoring overlapping and adjacent signals from genome-wide ChIP and DamID assays. Molecular BioSystems, 5, 1429.

Fu, Y. et al. (2008) The insulator binding protein CTCF positions 20 nucleosomes around its binding sites across the human genome. PLos Genet., 4, e1000138.

Gel, B. et al. (2015) regioneR: An r/bioconductor package for the association analysis of genomic regions based on permutation tests. Bioinformatics, btv562.

Heger, A. et al. (2013) GAT: A simulation framework for testing the association of genomic intervals. Bioinformatics, 29, 2046–2048.

Huen, D.S. and Russell, S. (2010) On the use of resampling tests for evaluating statistical significance of binding-site co-occurrence. BMC Bioinformatics, 11.

Johnson, W.E. et al. (2007) Adjusting batch effects in microarray expression data using empirical bayes methods. Biostatistics, 8, 118–27.

Kanduri, C. et al. (2018) Colocalization analyses of genomic elements: Approaches, recommendations and challenges. Bioinformatics, 35, 1615–1624.

Khushi, M. et al. (2014) Binding sites analyser (BiSA): Software for genomic binding sites archiving and overlap analysis. PLoS ONE, 9, e87301.

Layer, R.M. et al. (2018) GIGGLE: A search engine for large-scale integrated genome analysis. Nature Methods, 15, 123–126.

Moore, J.E. et al. (2020) Expanded encyclopaedias of DNA elements in the human and mouse genomes. 583, 699–710.

Nagraj, V. et al. (2018) LOLAweb: A containerized web server for interactive genomic locus overlap enrichment analysis. Nucleic Acids Research.

Quinlan, A.R. (2014) BEDTools: The swiss-army tool for genome feature analysis: BEDTools: The swiss-army tool for genome feature analysis. Current Protocols in Bioinformatics, 47, 11.12.1–11.12.34.

Sarmashghi, S. and Bafna, V. (2019) Computing the statistical significance of overlap between genome annotations with iStat. Cell Systems, 8, 523–529.e4.

Sheffield, N.C. and Bock, C. (2016) LOLA: Enrichment analysis for genomic region sets and regulatory elements in R and bioconductor. Bioinformatics, 32, 587–589.

Sheffield, N.C. and Furey, T.S. (2012) Identifying and characterizing regulatory sequences in the human genome with chromatin accessibility assays. Genes, 3, 651–70.

Simovski, B. et al. (2018) Coloc-stats: A unified web interface to perform colocalization analysis of genomic features. Nucleic Acids Research, 46, W186–W193.

Song, J. et al. (2011) Structure of DNMT1-DNA complex reveals a role for autoinhibition in maintenance DNA methylation. Science, 331, 1036–1040.

Thurman, R.E. et al. (2012) The accessible chromatin landscape of the human genome. Nature, 489, 75–82.

Wederell, E.D. et al. (2008) Global analysis of in vivo Foxa2-binding sites in mouse adult liver using massively parallel sequencing. Nucleic Acids Research, 36, 4549–4564.

Yu, G. et al. (2015) ChIPseeker: An r/bioconductor package for ChIP peak annotation, comparison and visualization. Bioinformatics, 31, 2382–2383.

Zhang, Z.D. et al. (2007) Statistical analysis of the genomic distribution and correlation of regulatory elements in the ENCODE regions. Genome Res., 17, 787–797.

Zhou, Y. et al. (2020) epiCOLOC: Integrating large-scale and context-dependent epigenomics features for comprehensive colocalization analysis. Frontiers in Genetics, 11.

